# Visualizing Effective Connectivity in the Human Brain

**DOI:** 10.1101/2025.03.06.641642

**Authors:** Alexander S. Atalay, Matteo Fecchio, Brian L. Edlow

**Affiliations:** Center for Neurotechnology and Neurorecovery, Department of Neurology, Massachusetts General Hospital and Harvard Medical School, Boston, MA, USA; Athinoula A. Martinos Center for Biomedical Imaging, Massachusetts General Hospital, Charlestown, MA, USA

## Abstract

To enhance mechanistic understanding of effective connectivity in the human brain, we created a tool that links high-temporal resolution transcranial magnetic stimulation electroencephalography (TMS-EEG) with high-spatial resolution diffusion MRI. This tool, Surface to Tractography Real-time EEG Activation Mapping in 4 Dimensions (STREAM-4D), integrates electrophysiologic source estimation models from TMS-evoked potentials (TEPs) with structural connectivity models from diffusion MRI tractography. In a proof-of-principle application, we used STREAM-4D to analyze TMS-EEG and diffusion MRI tractography data in a neurotypical subject across three stimulation sites: premotor, parietal, and occipital cortex. STREAM-4D revealed extensive structural connections associated with the TMS-evoked source models, involving thalamocortical, ipsilateral cortico-cortical, and transcallosal cortico-cortical connections. Activity-weighted structural connectivity differed for the three stimulation sites but shared two extensively connected nodes: the ipsilateral thalamus and putamen. STREAM-4D provides opportunities to elucidate how electrical activity evoked by a TMS pulse is related to underlying white matter architecture.

## Introduction

Effective connectivity, the causal influence of one brain region on others [1], is key to understanding brain function in normal and pathological conditions. Transcranial magnetic stimulation electroencephalography (TMS-EEG) has been used to assess effective connectivity in psychiatric and neurological disorders by perturbing cortical nodes within widely distributed brain networks [2-4]. However, the underlying structural properties that sustain the propagation of TMS-evoked potentials within these networks are not fully understood. This gap in knowledge is in part attributable to the lack of integrative tools that causally link structural brain networks to TMS-evoked potentials (TEPs).

To address this methodological gap and enhance mechanistic understanding of effective connectivity, we created a tool that links high-temporal resolution TMS-EEG with high-spatial resolution diffusion MRI: STREAM-4D (Surface to Tractography Real-time EEG Activation Mapping in 4 Dimensions). STREAM-4D integrates electrophysiologic source estimation models from TEPs with structural connectivity models from diffusion MRI tractography. This integration is assessed qualitatively using Blender, an open-source 3D animation suite, and quantitatively through an activation-weighted structural connectivity analysis in MRTrix [5]. By combining source estimation models and diffusion MRI tractography, STREAM-4D allows us to better understand how electrical activity evoked by a TMS pulse is related to underlying white matter architecture.

## Methods

We map source estimation to tractography by first extracting vertex and streamline endpoint coordinates from the cortical surface mesh and tractography data. Next, vertices are associated with streamline endpoints within a specified distance threshold using k-d tree spatial indexing. Finally, streamlines inherit the maximum of their parent surface vertices’ normalized source estimation intensity at each time point (**Figure 1**).

**Figure 1:**
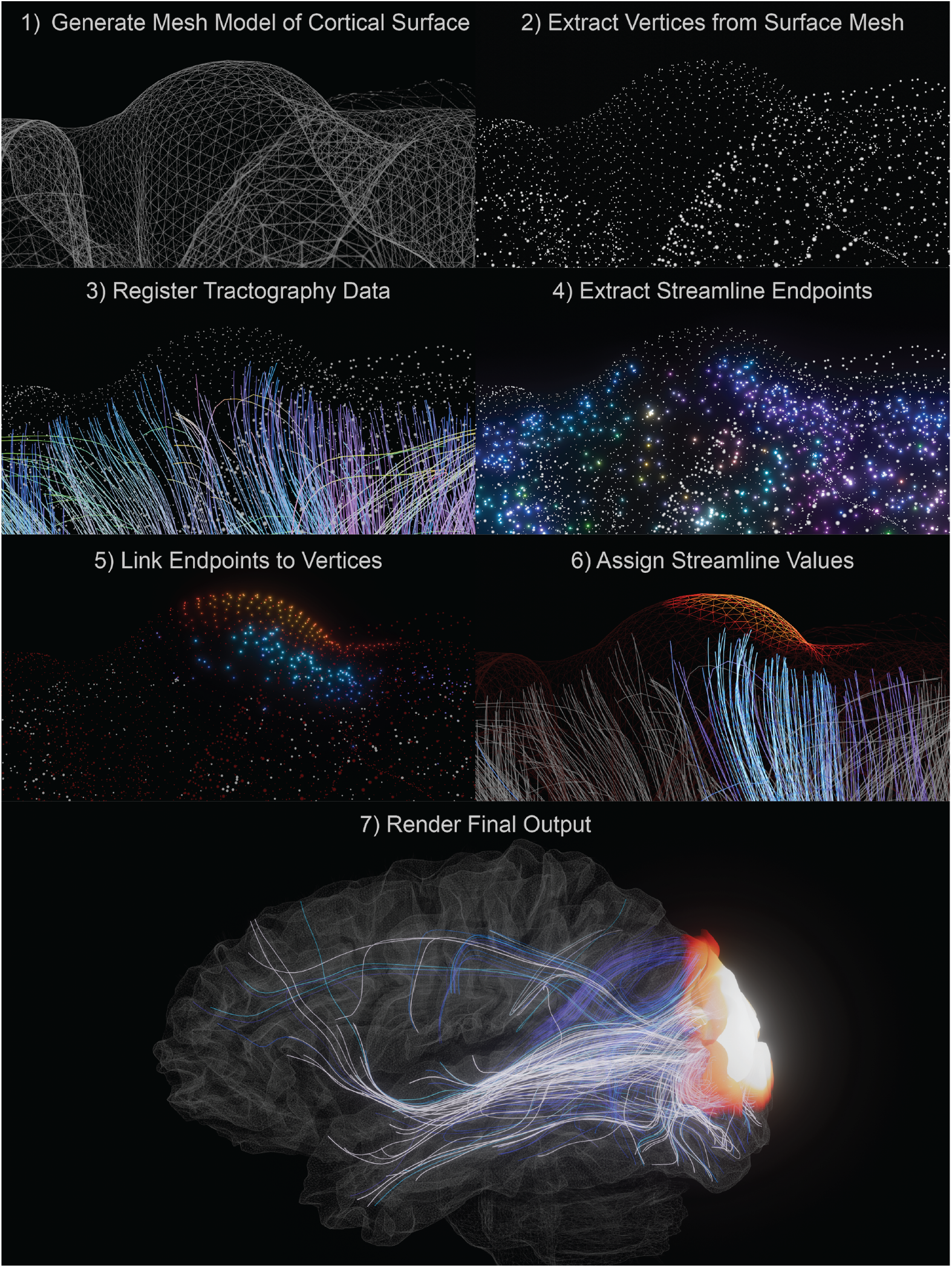
Overview of STREAM-4D Pipeline. Stepwise visualization of the STREAM-4D pipeline: 1) A cortical surface is generated and represented as a triangle mesh. 2) 3-dimensional coordinates for each surface vertex are extracted and saved to a coordinate matrix. 3) Streamlines are extracted from a tractography dataset, and 4) represented as directionally agnostic endpoints. 5) Streamline endpoints are linked to surface vertices using k-d tree spatial indexing. 6) Streamlines inherit the activation intensities of their parent vertices. 7) Time series data is animated and rendered in Blender for qualitative assessment.

To evaluate the associations qualitatively, a random subset of tractography streamlines is sampled from the activation matrix for rendering efficiency. Sampled streamlines are loaded into Blender as Non-Uniform Rational B-Splines curves and keyframed for color and emission using shader nodes and based on intensity values. The cortical surface is converted into a wavefront object, and the color and emission of each vertex are updated every frame based on the normalized source estimation scalars. To improve the clarity and interpretability of the visualization, only the top 10% of significant source activations are shown.

Quantitative evaluation is performed by first summing streamline activation intensity across the time series. Streamlines with temporally collapsed activation greater than 0 are extracted for structural connectivity analysis. Node regions are determined using FreeSurfer segmentations [6]. An MRTrix connectivity analysis is performed using ‘tck2connectome’ on extracted streamlines and weighted by the square of the activation array, multiplied by the output of ‘tcksift,’ which improves anatomical accuracy [5].

In a proof-of-principle application to the study of effective connectivity, we used STREAM-4D to analyze TMS-EEG and diffusion MRI tractography data in a 44-year-old neurotypical man. The subject provided informed consent to participate in a research protocol approved by the Mass General Brigham Institutional Review Board [7]. He was scanned with an MRI protocol that included a T1-weighted multi-echo magnetization-prepared rapid gradient echo sequence for surface mapping and a diffusion-weighted sequence for tractography. Diffusion data were acquired with a spin-echo echo-planar imaging sequence (TR=3,230 ms, TE=89 ms) with b-values of 0, 1,500, and 3,000 s/mm^2^, 100 diffusion-encoding directions, a 1.5 mm isotropic voxel size, and opposing phase-encoding directions (anterior-posterior and posterior-anterior) to correct susceptibility-induced distortions.

After MRI acquisition, the subject underwent TMS over premotor, parietal, and occipital cortices (Nexstim NBT 2.2, Nexstim, Finland), with simultaneous recording of evoked EEG responses via a 64-electrode cap (EASYCap) connected to BrainAmp DC amplifiers (Brain Products GmbH, Germany). We anatomically identified cortical targets on the T1-weighted MRI co-registered with the subject’s head. The position of EEG electrodes with respect to the subject’s scalp was digitized, stored in the neuronavigation system, and subsequently used to compute the forward model for source estimation. TEP acquisition and EEG processing steps are detailed in Supplementary Methods.

Following the preprocessing of diffusion MRI data, we created basis functions for each tissue type, constructed a grey-white matter boundary for seed analysis, and generated streamlines using MRTrix3 [8]. Using the mne python package [9], we created a Boundary Element Method conductivity model and generated a source space with ico5 spacing. Next, we made a forward model, calculated a covariance matrix using the ‘empirical’ method, and obtained an inverse solution using a minimum norm estimate model [10]. For each source, the prestimulus period was used to establish a baseline activity distribution and significant activation was determined using a Z-score threshold (p<0.05) and corrected for multiple comparisons using the Bonferroni method, treating each source as an independent metric. Finally, we integrated the tractography output and significant source activation using STREAM-4D and a vertex-to-endpoint distance threshold of 3 mm.

## Results

At each of the three stimulation sites – premotor, parietal, and occipital cortex – STREAM-4D revealed structural connections activated by the TMS-evoked electrical waves involving thalamocortical, ipsilateral cortico-cortical, and transcallosal cortico-cortical connections (**Figure 2**; **Video S1**). Activity-weighted structural connectivity differed for the three stimulation sites but shared two extensively connected nodes: the ipsilateral thalamus and putamen (**Figure 3**).

**Figure 2:**
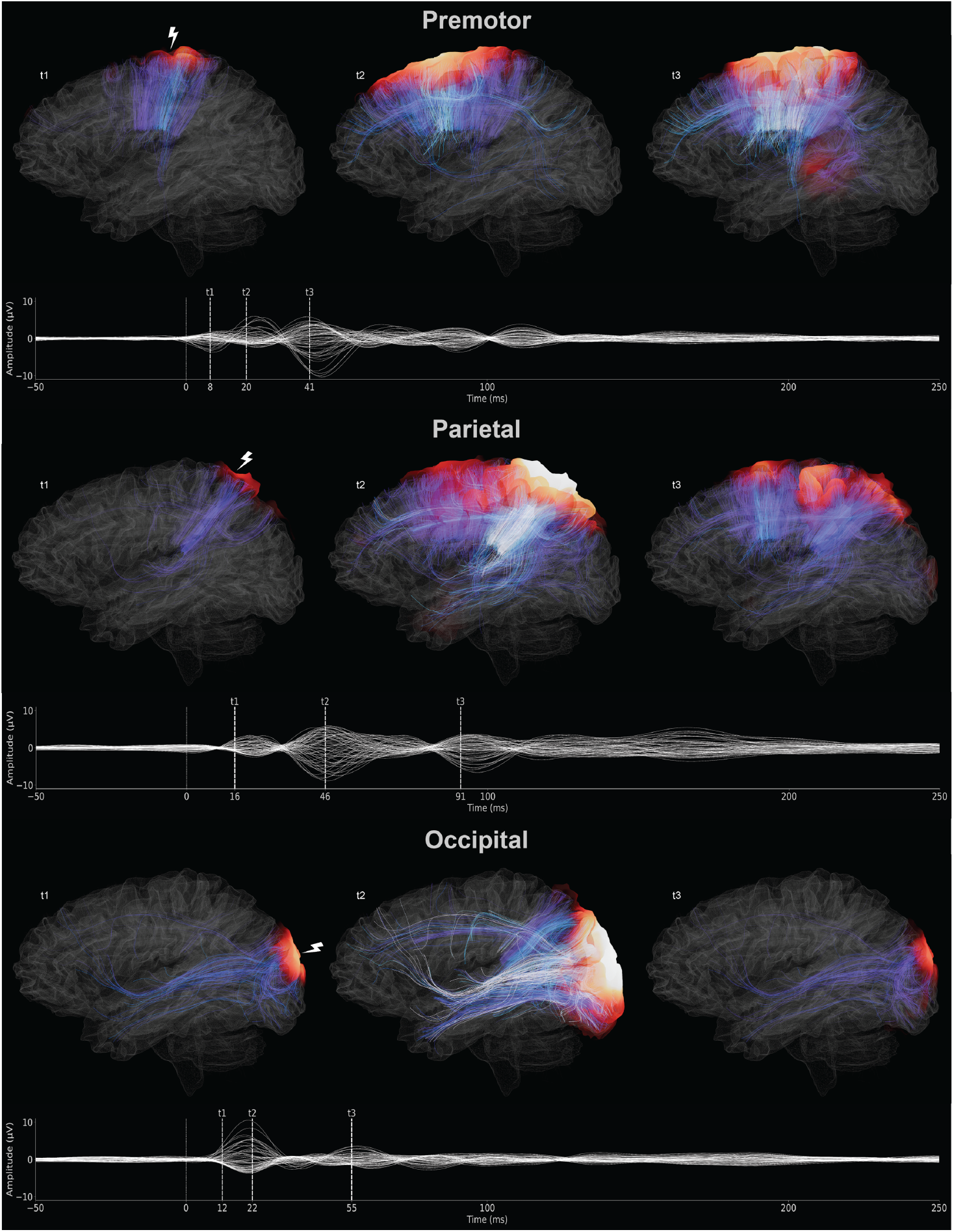
Qualitative Assessment of STREAM-4D Across Stimulation Sites. Streamline and surface activation time series data are plotted using Blender for each stimulation site: premotor, parietal, and occipital. Snapshots are shown at three electrophysiologic event time points (t1, t2, and t3) and displayed above butterfly plots showing each EEG channel’s evoked potential across the stimulation event (t=0). Streamline and surface activation intensity are indicated by emission color and opacity; a “hot” colormap is used to show surface activation intensity, and a “cool” colormap is used to show the activation intensity of associated streamlines. Lightning bolt glyphs are used to indicate the cortical location of each stimulation site. Each stimulation site reveals extensive thalamocortical, ipsilateral and transcallosal cortico-cortical structural connections shown by the activated streamlines that may support the TEP activations.

**Figure 3:**
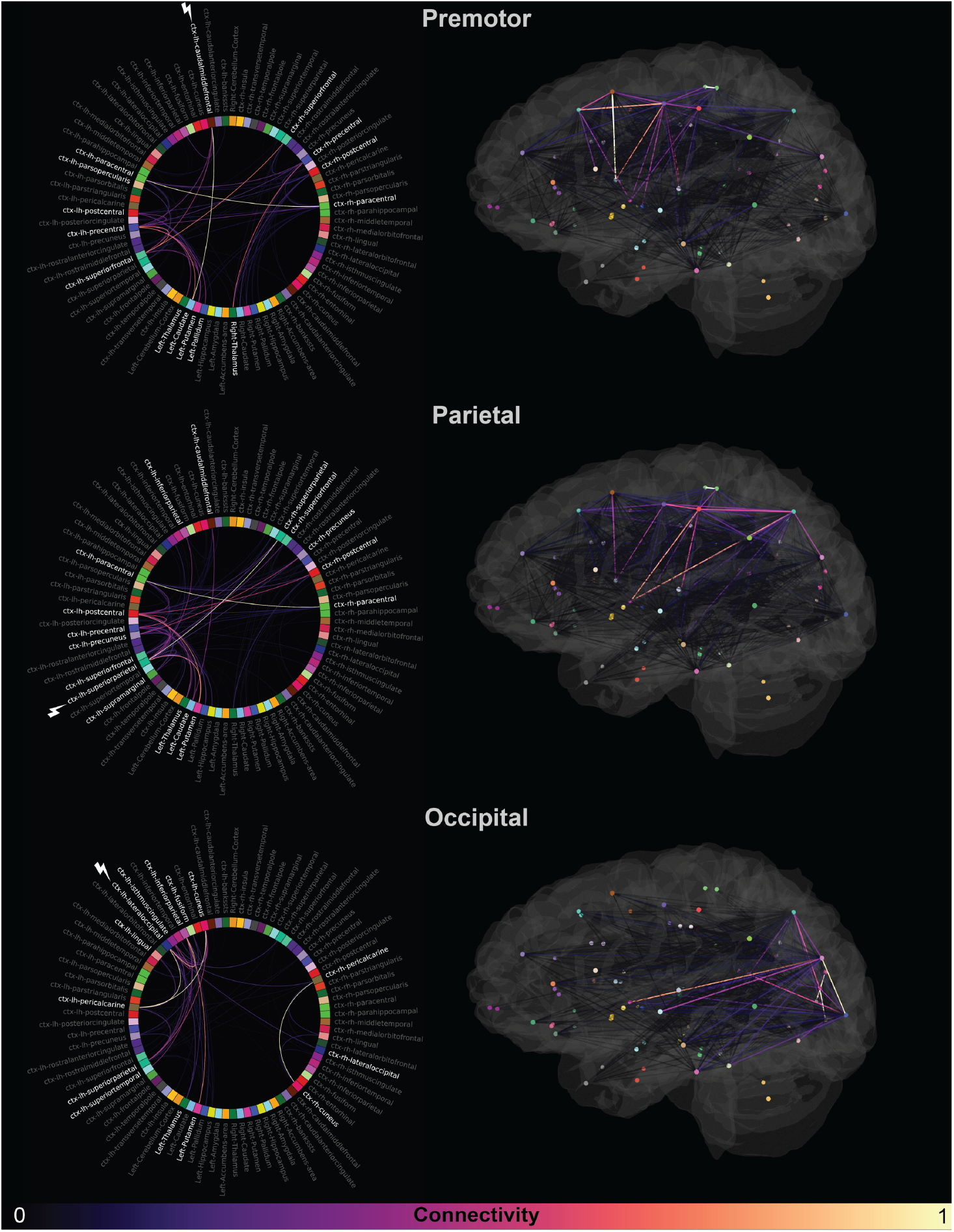
Activity-Weighted Structural Connectivity of Stimulation Sites. 2D connectograms and 3D structural network visualizations of activity-weighted structural connectivity for each stimulation site (premotor, parietal, and occipital). A structural connectivity analysis was performed on the streamlines associated with each stimulation and weighted by integrated activation. Connectivity nodes are created using the center of mass of FreeSurfer segmentations and colored using the default FreeSurfer “aseg” lookup table. The color of edges between nodes represents normalized connectivity. To improve interpretability, labels of the nodes that demonstrate connectivity in the top 5% of each stimulation site are white; all other nodes are grey. A lightning bolt glyph indicates the node spatially corresponding to the stimulation site. The ipsilateral thalamus and putamen are the only network nodes in the top 5% of activity-weighted structural connectivity for all three stimulation sites.

While larger studies are needed to clarify the relative contributions of thalamocortical and corticocortical networks to the human brain’s physiologic response to a TMS pulse, these proof-of-principle STREAM-4D results reveal previously unseen relationships between structural networks and TMS-EEG measurements of effective connectivity. Systematic application of STREAM-4D to multimodal TMS-EEG-neuroimaging studies in patients with neuropsychiatric disorders has the potential to shed mechanistic light on the anatomic basis of effective connectivity in the human brain.

We distribute the STREAM-4D code to facilitate the study of effective connectivity (https://github.com/ComaRecoveryLab/STREAM-4D). The code is adaptable and can support integrative, multimodal analysis of TMS-EEG data with multiple types of neuroimaging data. Hence, we envision that STREAM-4D can be used to advance the study of structure-function relationships in the human brain.

## Supplementary data

All figures and the supplementary video can be found online at https://zenodo.org/records/14974861. The supplementary video can also be viewed online at https://youtu.be/V0hqs-noYNg.

## References

[1] Friston KJ. Functional and eBective connectivity: a review. Brain Connect 2011;1(1):13–36.

[2] Casali AG, Gosseries O, Rosanova M, Boly M, Sarasso S, Casali KR, et al. A theoretically based index of consciousness independent of sensory processing and behavior. Sci Transl Med 2013;5(198):198ra05.

[3] Tremblay S, Rogasch NC, Premoli I, Blumberger DM, Casarotto S, Chen R, et al. Clinical utility and prospective of TMS-EEG. Clin Neurophysiol 2019;130(5):802–44.

[4] Esposito R, Bortoletto M, Zaca D, Avesani P, Miniussi C. An integrated TMS-EEG and MRI approach to explore the interregional connectivity of the default mode network. Brain Struct Funct 2022;227(3):1133–44.

[5] Smith RE, Tournier JD, Calamante F, Connelly A. SIFT2: Enabling dense quantitative assessment of brain white matter connectivity using streamlines tractography. NeuroImage 2015;119:338–51.

[6] Fischl B. FreeSurfer. NeuroImage 2012;62(2):774–81.

[7] Edlow BL, Fecchio M, Bodien YG, Comanducci A, Rosanova M, Casarotto S, et al. Measuring Consciousness in the Intensive Care Unit. Neurocrit Care 2023;38(3):584–90.

[8] Christiaens D, Reisert M, Dhollander T, Sunaert S, Suetens P, Maes F. Global tractography of multi-shell diBusion-weighted imaging data using a multi-tissue model. NeuroImage 2015;123:89–101.

[9] Gramfort A, Luessi M, Larson E, Engemann DA, Strohmeier D, Brodbeck C, et al. MEG and EEG data analysis with MNE-Python. Front Neurosci 2013;7:267.

[10] Hamalainen MS, Ilmoniemi RJ. Interpreting magnetic fields of the brain: minimum norm estimates. Med Biol Eng Comput 1994;32(1):35–42.

