## Supplementary Methods for "Visualizing Effective Connectivity in the Human Brain"

#### TMS-EEG data acquisition and processing

TMS stimulation (Nexstim NBT 2.2, Nexstim, Finland) with simultaneous recording of evoked EEG responses (64-electrode cap EASYCap connected to BrainAmp DC amplifiers, Brain Products GmbH, Germany) was performed by targeting premotor, parietal, and occipital cortexes. For each cortical target 220 biphasic pulses were delivered at an inter-pulse interval randomly jittered between 1-1.3 s. At the same time, a continuous masking noise was played throughout the EEG recordings to prevent auditory contaminations due to TMS discharge [1]. Stimulation parameters were optimized at each cortical target to elicit TMS-evoked potentials characterized by peak-to-peak amplitude in the early (8–50 ms) components in the EEG channels located underneath the TMS coil of at least 10 μV [2]. Thus, stimulation intensity was set at 51, 68, and 50 percent of the maximal output (128, 127, 117 V/m intensity of the electric field induced by TMS at the cortical target), respectively, for the premotor, parietal, and occipital cortexes. The position of EEG electrodes with respect to the subject's scalp was digitized and stored in the neuronavigation system at the end of the experimental session.

EEG data were visually inspected to discard epochs with discontinuities and bad channels. At least 152 good epochs were selected for each session and no EEG channels were deemed as bad and thus removed. TMS magnetic artifacts were removed by replacing the interval around the TMS pulse (-2 to 5 ms) with the preceding interval (-9 to -2 ms) and by applying a moving-average filter (5th order) between 3 and 7 ms. Prior to data epoching (1600 ms around the TMS pulse), a high-pass filter at .01 Hz (1st order, IIR) and a 1 Hz high-pass filter (3rd order, Butterworth) were applied to remove DC fluctuations, while a bandstop filter at 60 Hz (3rd order, Butterworth) was employed to remove line noise. Then, data were referenced to the average of the signal recorded at all EEG electrodes, and the ICA was performed to remove eye movement artifacts and spontaneous muscle activity. Finally, data were low-pass filtered at 45 Hz (3rd order, Butterworth), downsampled to 1 kHz, and additionally segmented from -600 to 600 ms around the TMS pulse.

#### Diffusion MRI preprocessing

Diffusion MRI (dMRI) data were preprocessed using a standard pipeline consistent with prior protocols. dMRI data were corrected for thermal noise using the 'dwidenoise' command from MRTrix3 [3]. Gibbs deranging was performed using the 'dwidegibbs' MRTrix3 function. Correction of susceptibility-induced (EPI) distortions, eddy currents, and subject motion was performed using the FSL commands 'topup' and 'eddy' [4].

#### Supplementary Figures:

High resolution versions of all figures can be accessed at the following link: https://zenodo.org/records/14974861.

#### Video S1: Overview of STREAM-4D Pipeline

Streamline and surface activation time series are rendered using Blender for each stimulation site: premotor, parietal, and occipital. Corresponding butterfly plots show each

EEG channel's evoked potential relative to the stimulation event (dashed vertical line). Streamline and surface activation intensity are indicated by emission color and opacity; a “hot” colormap is used to show surface activation intensity, and a “cool” colormap is used to show the activation intensity of associated streamlines. Each stimulation site reveals extensive thalamocortical, ipsilateral and transcallosal cortico-cortical structural connections shown by the activated streamlines that may support the TEP perturbations. The video can be viewed at https://youtu.be/V0hqs-noYNg.

### Supplementary References


[1] Russo S, Sarasso S, Puglisi GE, Dal Palu D, Pigorini A, Casarotto S, et al. TAAC - TMS Adaptable Auditory Control: A universal tool to mask TMS clicks. J Neurosci Methods 2022;370:109491.

[2] Casarotto S, Fecchio M, Rosanova M, Varone G, D'Ambrosio S, Sarasso S, et al. The rt-TEP tool: real-time visualization of TMS-Evoked Potentials to maximize cortical activation and minimize artifacts. J Neurosci Methods 2022;370:109486.

[3] Tournier JD, Smith R, Raffelt D, Tabbara R, Dhollander T, Pietsch M, et al. MRtrix3: A fast, flexible and open software framework for medical image processing and visualisation. NeuroImage 2019;202:116137.

[4] Andersson JL, Skare S, Ashburner J. How to correct susceptibility distortions in spin-echo echo-planar images: application to diffusion tensor imaging. NeuroImage 2003;20(2):870-88.
